# The genetics of Bene Israel from India reveals both substantial Jewish and Indian ancestry

**DOI:** 10.1101/025809

**Authors:** Yedael Y. Waldman, Arjun Biddanda, Natalie R. Davidson, Paul Billing-Ross, Maya Dubrovsky, Christopher L. Campbell, Carole Oddoux, Eitan Friedman, Gil Atzmon, Eran Halperin, Harry Ostrer, Alon Keinan

**Affiliations:** Department of Biological Statistics and Computational Biology, Cornell University, Ithaca, NY 14853, USA; Department of Molecular Microbiology and Biotechnology, Tel Aviv University, Ramat Aviv, Tel Aviv 6997801, Israel; Danek Gertner Institute of Human Genetics, Chaim Sheba Medical Center, Tel-Hashomer 52621, Israel; Sackler School of Medicine, Tel Aviv University, Ramat Aviv, Tel Aviv 6997801, Israel; Department of Pathology, Albert Einstein College of Medicine, Bronx, NY 10461,USA; Departments of Medicine and Genetics, Albert Einstein College of Medicine, Bronx, NY 10461, USA; Department of Human Biology, Faculty of Natural Sciences, University of Haifa, Haifa, Israel.; The Blavatnik School of Computer Science, Tel Aviv University, Ramat Aviv, Tel Aviv 6997801, Israel; International Computer Science Institute, Berkeley, California 94704, USA; Department of Pediatrics, Albert Einstein College of Medicine, Bronx, NY 10461,USA; These authors contributed equally to this work; Current address: Computational Biology Center, Memorial Sloan Kettering Cancer Center, New York, NY 10044, USA

**Author notes:** Corresponding author: Alon Keinan, Department of Biological Statistics and Computational Biology, Cornell University, Ithaca, NY 14853, USA.

## Abstract

The Bene Israel Jewish community from West India is a unique population whose history before the 18^th^ century remains largely unknown. Bene Israel members consider themselves as descendants of Jews, yet the identity of Jewish ancestors and their arrival time to India are unknown, with speculations on arrival time varying between the 8th century BCE and the 6th century CE. Here, we characterize the genetic history of Bene Israel by collecting and genotyping 18 Bene Israel individuals. Combining with 486 individuals from 41 other Jewish, Indian and Pakistani populations, and additional individuals from worldwide populations, we conducted comprehensive genome-wide analyses based on F_ST_, principal component analysis, ADMIXTURE, identity-by-descent sharing, admixture linkage disequilibrium decay, haplotype sharing and allele sharing autocorrelation decay, as well as contrasted patterns between the X chromosome and the autosomes. The genetics of Bene Israel individuals resemble local Indian populations, while at the same time constituting a clearly separated and unique population in India. They are unique among Indian and Pakistani populations we analyzed in sharing considerable genetic ancestry with other Jewish populations. Putting together the results from all analyses point to Bene Israel being an admixed population with both Jewish and Indian ancestry, with the genetic contribution of each of these ancestral populations being substantial. The admixture took place in the last millennium, about 19-33 generations ago. It involved Middle-Eastern Jews and was sex-biased, with more male Jewish and local female contribution. It was followed by a population bottleneck and high endogamy, which can lead to increased prevalence of recessive diseases in this population. This study provides an example of how genetic analysis advances our knowledge of human history in cases where other disciplines lack the relevant data to do so.

## Introduction

How well does the oral history of a group reflect its origins? The Bene Israel community in West India is a unique community whose historical background before the 18^th^ century other than their oral history remains largely unknown [1-3]. The Jewish philosopher, Maimonides, in a letter written 800 years ago (circa 1200 CE), briefly mentioned a Jewish community living in India and may have referred to them [4]. In the 18^th^ century Bene Israel members lived in villages along the Indian Konkan coast and were called *Shanivar Teli* (Marathi for ‘Saturday oil pressers’), as they were oil pressers who did not work on Saturdays. After 1948, most of the community immigrated to Israel. At the beginning of the 21^st^ century, approximately 50,000 members lived in Israel, whereas about 5,000 remained in India, mainly in Mumbai [2]. Oral history among Bene Israel holds that they are descendants of Jews whose ship wrecked on the Konkan shore, with only seven men and seven women surviving [2, 3, 5]. The exact timing of this event, as well as the origin and identity of the survivors, are not part of this oral history. Some date it around two millennia ago [2], whereas others suggest a specific date and origin: around 175 BCE, where the survivors were Jews living in the northern parts of the land of Israel that left their homes during the persecutions of Antiochus Epiphanes [5]. Adding to the vagueness of Bene Israel origin is the fact that a similar story of seven surviving couples is found in the oral histories of other Indian populations [2, 3]. Others suggest that the ancestors of Bene Israel arrived to India earlier – as early as the 8^th^ century BCE – or later – from Yemen, during the first millennium CE or from Southern Arabia or Persia in the 5^th^ or 6^th^ century CE [4]. However, beyond vague oral history and speculations, there has been no independent support for any of these claims, and Bene Israel origin and whether they are related at all to other Jewish populations and remain “shrouded in legend” [4].

In the last decades, genetic information has become an important source for the study of human history and has been applied numerous times for various Jewish populations, first based on uniparental Y chromosomal and mitochondrial DNA (mtDNA) markers [6-11] and later by using genome-wide markers [12-15]. These studies found that most Jewish Diasporas share ancestry that can be traced back to the Middle-East, in accordance with historical records [12-15]. Some of these studies included Bene Israel members, though with inconclusive results [15]. Bene Israel’s mtDNA pool was shown to consist of mostly local Indian origin [8, 11, 13] although a few haplogroups found in Bene Israel samples were not present in local Indian populations, but were present in several Jewish populations [11]. A Y chromosome analysis hinted at paternal link between Bene Israel and the Levant, but the study was based on only four Bene Israel males [13]. Another Y chromosome analysis showed that a common Indian haplogroup was almost absent in Bene Israel males, whereas the Cohen Modal Haplotype (CMH) [16] was common in Bene Israel (and other Jewish) males though also present at lower frequencies in other Indian populations [3]. These results suggest that the founding males of this population might have had Middle Eastern, possibly Jewish, origins. On the contrary, analysis of the autosomes or the X chromosome did not find any evidence of Jewish origin of Bene Israel, and it has been concluded that they resembled other Indian populations [13, 15]. Thus, genetic studies to date left the genetic history of Bene Israel largely unknown.

The complex genetic structure of Indian populations imposes a great challenge for genetic analysis of Bene Israel. Previous studies showed that most contemporary Indian populations are a result of ancient admixture (64-144 generations ago) of two genetically divergent populations: Ancestral North Indians (ANI), who are related to west Eurasians, and Ancestral South Indians (ASI), who are not closely related to populations outside India and related to indigenous Andaman Island people [17, 18]. Different Indian populations vary in the proportion of admixture between these two ancestral populations [18]. As the ANI component is related to west Eurasia, which includes the Middle East and Europe, a study analyzing the connection between Bene Israel and other Jewish or Middle Eastern populations needs to examine whether such a connection reflects a unique ancestry component, rather than simply being a result of the large ANI component.

To study the genetic history of Bene Israel, while addressing this challenge, we present here the largest collection of Bene Israel individuals that has been assayed genome-wide to date (18 individuals), and we use the collection in conjunction with genotype data of 486 individuals from 41 other Jewish, Indian and Pakistani populations, as well as samples from various worldwide populations. We apply an array of genome-wide population genetics tools to characterize the origins of Bene Israel and their relations to both Indian and Jewish populations, uncovering the genetic history of this unique population.

## Results

### Bene Israel cluster with Indian populations but as a distinct group

We genotyped Bene Israel individuals and combined the data with 14 other Jewish populations from worldwide Diaspora previously genotyped using the same array [12, 14]. We applied various quality control (QC) steps on these samples, resulting with 18 individuals of the Bene Israel community together with 347 samples from the other 14 Jewish populations. We also applied the same QC steps to a different dataset with samples from 18 different Indian populations (96 individuals) [17], as well as HapMap3 populations, that were genotyped previously on the same array [17] and merged the two datasets. We also merged the data with additional populations from the HGDP panel [19]: three non-Jewish Middle Eastern populations (Druze, Bedouin and Palestinians), and nine Pakistani populations (Table S1). The Middle-Eastern populations were used to distinguish between Middle-Eastern and Jewish specific ancestry while the Pakistani populations were used to represent populations that are geographically located between India and the Middle-East. In addition, some Pakistani populations are also part of the ANI-ASI admixture, with a relatively large ANI component as compared to the Indian populations [17, 18] and therefore also represent this ancient admixture. As the merging with HGDP resulted in considerable reduction in number of SNPs available for analysis, we only considered this merged dataset for some analyses.

PCA (Principal Component Analysis) of the merged dataset, including four HapMap populations (YRI, CEU, CHB and JPT; total 873 individuals) showed that Jewish populations cluster together with Europeans and Middle-Eastern populations, while Indian and Pakistani populations form their own cluster, between East-Asians and Jews/Europeans (Figure 1A). The Bene Israel population clustered with the Indian and Pakistani populations, similar to the results of a previous study [13]. Bene Israel was the closest to the Jewish/Middle-Eastern/European cluster as compared to all other Indian populations, while several Pakistani populations were even closer to that cluster than Bene Israel (Figure 1A). When focusing only on Jewish, Middle-Eastern, Pakistani and Indian samples (Figure 1B), the first PC separated between Indian/Pakistani and Jewish/Middle-Eastern populations while the second PC spanned the Jewish populations. Members of non-Jewish Middle-Eastern populations were located within the Jewish cluster, near Middle-Eastern Jewish populations. Indian populations were ordered based on their ANI-ASI admixture [17, 18] such that populations with higher ANI proportion were closer to Jews in general, as expected from the Middle-Eastern origins of the latter [12-15]. Bene Israel members were located closely to members of other Indian populations but were also the closest to samples from Jewish populations among all Indian samples, while some Pakistani populations were more similar to Jewish and Middle-Eastern populations (Figure 1B). PCA with only Bene Israel, Indian and Pakistani population showed the ANI-ASI incline spanning both Indian and Pakistani populations, while Bene Israel were near populations with high ANI component, but slightly off the incline, perhaps suggesting a different origin. (Figure 1C). PCA of only Bene Israel, Pakistani, Middle-Eastern and Jewish populations showed the separation of Bene Israel members from other Jewish populations and the fact that some Pakistani populations were closer than Bene Israel to Jewish populations (Figure 1D). To avoid bias in PCA due to differences in number of samples between populations [20], we repeated the analysis while limiting the number of samples from each population to 4, and obtained similar results (Figure S1).

**Figure 1.**
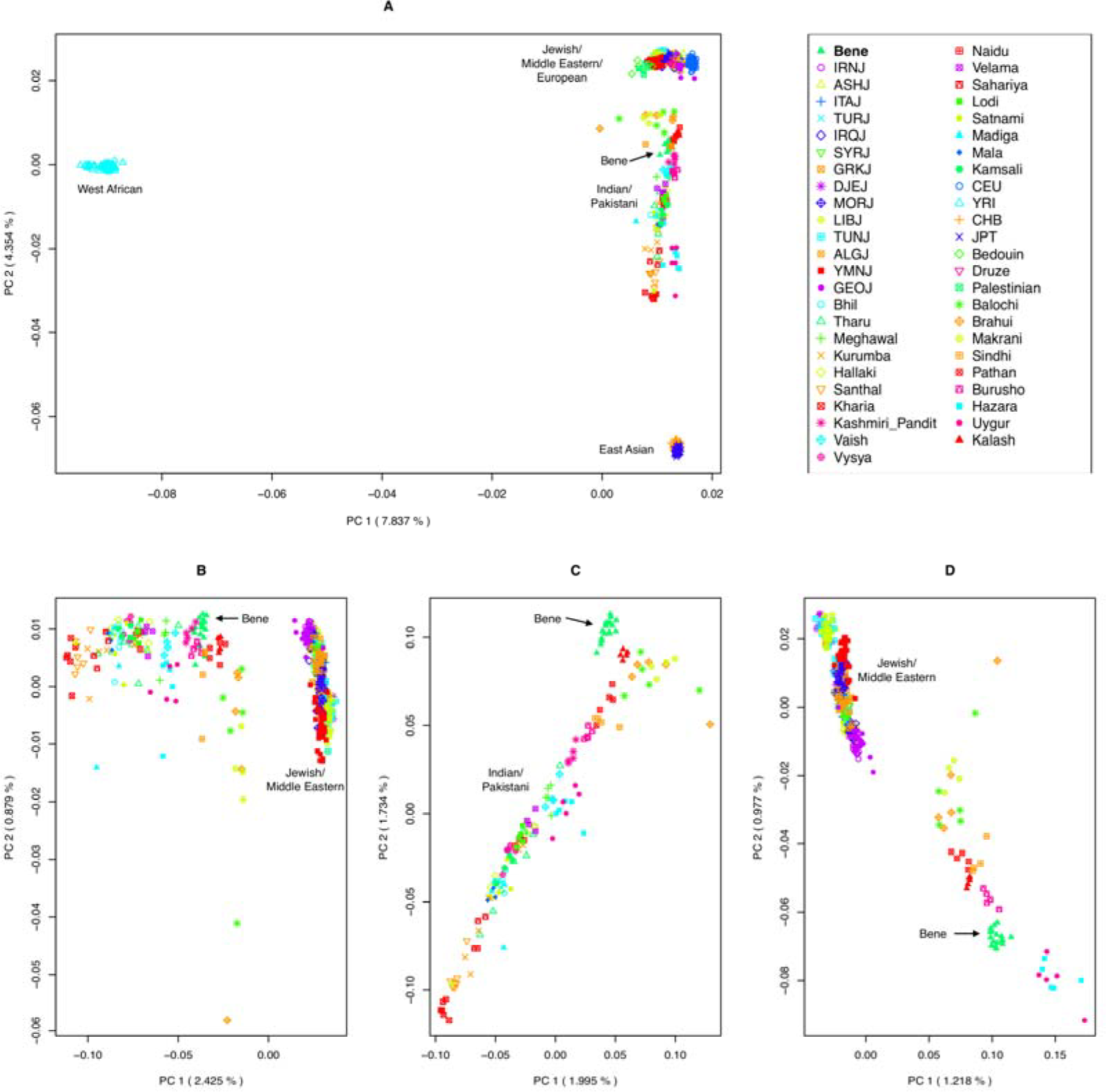
Principal Component Analysis of Jewish, Indian and worldwide populations. Each panel presents the top two principal components for a set of populations that include Bene Israel together with: (A) Jewish, Indian, Pakistani, Middle Eastern and four worldwide HapMap populations (CEU, CHB, JPT and YRI); (B) Jewish, Middle-Eastern, Pakistani and Indian populations; (C) Indian and Pakistani populations; (D) Jewish, Middle-Eastern and Pakistani populations. Abbreviations of Jewish populations: Bene Israel (Bene), Algerian Jews (ALGJ), Ashkenazi Jews (ASHJ), Djerban Jews (DJEJ), Georgian Jews (GEOJ), Greek Jews (GRKJ), Iranian Jews (IRNJ), Iraqi Jews (IRQJ), Italian Jews (ITAJ), Libyan Jews (LIBJ), Moroccan Jews (MORJ), Syrian Jews (SYRJ), Tunisian Jews (TUNJ), Yemenite Jews (YMNJ).

We also examined the relation between Bene Israel and other Indian and Jewish populations using the F_ST_ statistic, which measures genetic drift between populations based on differences in allele frequencies [17, 21] (Figure S2 and Table S2). This analysis revealed the isolation and genetic drift of Bene Israel from both Indian and Jewish populations: While the average F_ST_ between pairs of Jewish populations was 0.011, the average F_ST_ between Bene Israel and other Jewish populations was significantly higher (0.04, Wilcoxon rank sum P-value=1.97e-9). Similarly, while the average F_ST_ between different Indian populations was 0.011, the mean F_ST_ between Bene Israel and other Indian populations was significantly higher (0.033, P-value=8.63e-12, Wilcoxon rank sum test).

### ADMIXTURE analysis suggests Bene Israel members have Middle-Eastern ancestry

ADMIXTURE [22] assigns for each individual its proportion in any of a set of hypothetical ancestral populations and hence can reveal relations between different populations. We used this tool on our dataset for varying values of K (the number of hypothetical ancestral populations) on the same set of 873 individuals from the first PCA (Figure 2 and Figure S3). For K=3, we observed three clusters: East Asian, Sub-Saharan African and Middle Eastern/European. Indian and Pakistani populations were mainly composed from Middle-Eastern/European and East-Asian components. The proportion of the Middle-Eastern/European component in the Indian and Pakistani populations was highly correlated to their ANI component in the ANI-ASI admixture [17] (R=0.98, P-value<e-16, Spearman correlation). Among the Indian populations, Bene Israel had the highest proportion of Middle-Eastern/European component, but it was comparable to that of some other Pakistani populations. At K=4 an Indian cluster emerged which reflects the ASI component in these populations and most Indian and Pakistani populations were composed from this component and the Middle-Eastern/European component, while some of them also had an East-Asian component. Again, Bene Israel and three Pakistani populations (Balochi, Brahui and Makrani) had the highest proportions of Middle-Eastern/European component among Indian and Pakistani populations. At K=5, the European/Middle-Eastern cluster was divided into two clusters: European (reflected by the European population CEU) and Middle-Eastern (reflected by Jewish and Middle-Eastern populations). Importantly, Bene Israel population exhibited a different trend as compared to other Indian populations: While the ANI component of Indian populations was now mainly reflected in the European component, Bene Israel showed a significantly higher proportion of a Middle-Eastern component (mean 29.5% as compared to less than 14% in all members of other Indian populations, Wilcoxon rank sum P-value=1.75e-12). Nevertheless, some Pakistani populations showed similar or even higher proportions of the Middle-Eastern component (e.g., mean of 33% for Makrani). At K=6, which provided the best fit based on cross-validation, a new cluster emerged which was mainly found in North-African Jews (Djerban, Libyan and Tunisian Jews). At K=7 a new cluster represented Iranian Jews, but was also present in larger fractions in other Middle-Eastern and Asian populations, including Bene Israel. Interestingly, at K=8 Bene Israel formed their own hypothetical ancestral component, marking again the uniqueness of this population and its deviation from other populations. This ancestral component was present, in minute proportions, in many Indian and Pakistani populations but also in some Middle-Eastern populations (Jewish and non-Jewish).

**Figure 2.**
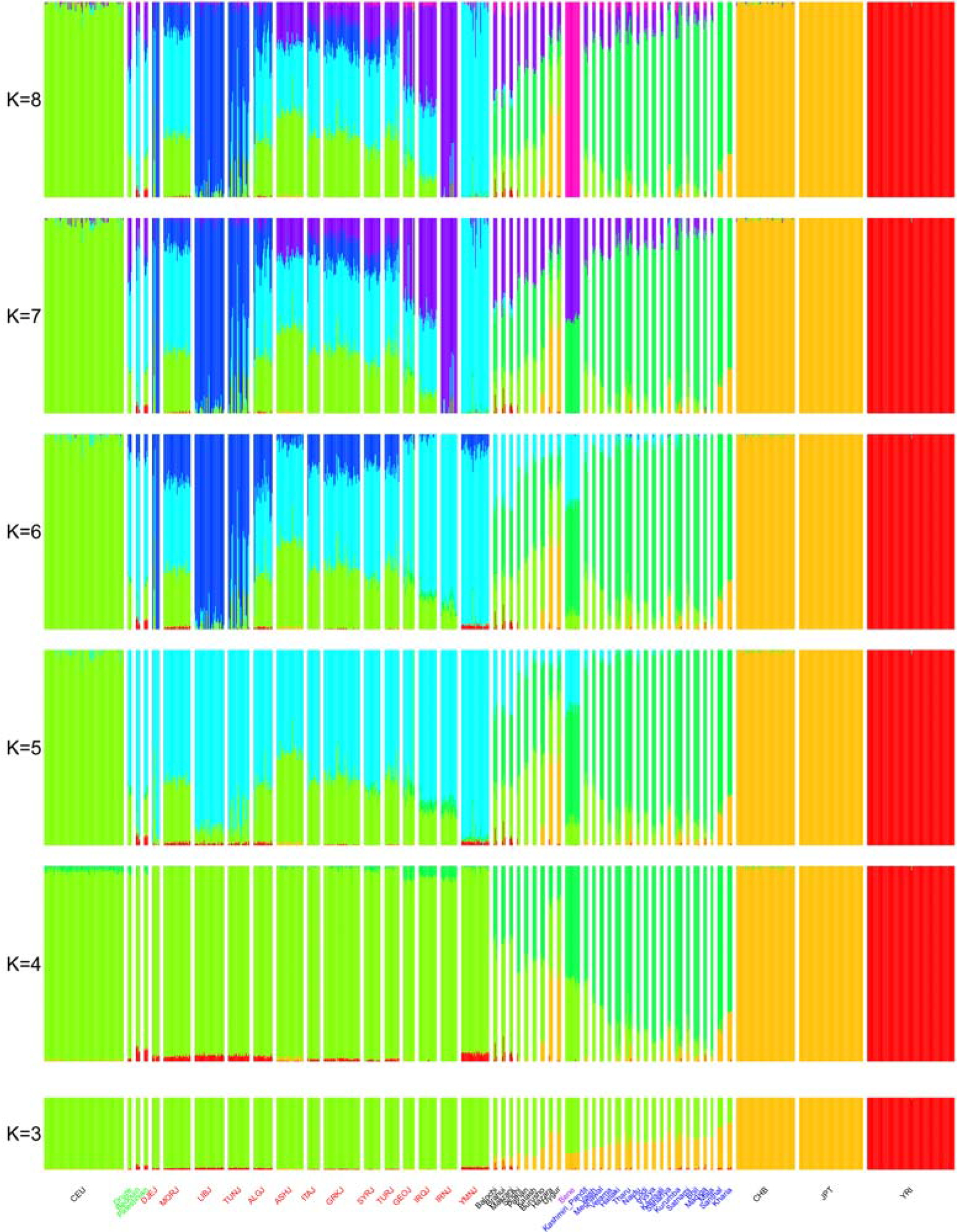
ADMIXTURE analysis for Jewish, Indian, Pakistani, Middle Eastern (Druze, Bedouin and Palestinians) and representative HapMap (CEU, YRI, JPT and CHB) populations. K, the number of clusters, varies from K=3 to K=8. We colored some of the populations names based on the following groups: Bene Israel (purple), Jews (red), Indian (blue) and Middle-Eastern (green). See also Figure S3.

### Identity-by-descent analysis suggest Bene Israel members are related to Jewish populations

The results based on PCA and ADMIXTURE, show that Bene Israel is more closely-related to Middle-Eastern and Jewish populations as compared to all other Indian populations examined here. However, the same claim cannot be made when compared to some Pakistani populations. Therefore, the question remains: Does Bene Israel have Jewish or Middle-Eastern ancestry that is not shared by other Asian populations? Next, we analyzed the relations between populations based on Identity-by-descent (IBD) sharing of their individuals. IBD segments shared by two individuals represent a segment inherited from a common ancestor. Higher IBD sharing, and specifically of long segments, suggests a more recent common ancestor without intervening recombination [23]. Following a similar previous analysis of the Jewish populations examined here (except Bene Israel) [12, 14], we used GERMLINE [24] to detect IBD segments between individuals and defined the IBD sharing between individuals to be the total length (in cM units) of IBD segments shared between the two individuals. IBD sharing between populations was defined as the average IBD sharing of unrelated individuals from these populations. As expected from previous results, each Jewish population exhibited significant higher IBD sharing with other Jewish populations than with Indian populations and each Indian population exhibited higher IBD sharing with other Indian populations than with Jewish populations (Wilcoxon P-value<0.05 for all populations; Figure 3A). Having these two IBD clusters of Indian and Jewish populations, we observed that compared to all Jewish populations, Bene Israel had the highest IBD sharing with all Indian populations, and compared to all Indian populations, Bene Israel had the highest IBD sharing with all Jewish populations (Figure S4A). Furthermore, the only population with no significant IBD sharing between the two clusters of Jewish and Indian populations was Bene Israel: (mean IBD sharing=18.24 cM vs. 18.19 cM with Indian and Jewish populations, respectively. P-value=0.61; Wilcoxon rank sum test; Figure 3B). Middle-Eastern Jews (specifically Georgian, Iraqi, Syrian and Iranian Jews) showed significantly higher average IBD sharing as compared to all other Jewish populations examined (Wilcoxon test P-value <e-10 for all pairs). Interestingly, the closest Indian populations to Bene Israel (Velama, Lodi and Bhil) were not those with the highest ANI component.

**Figure 3.**
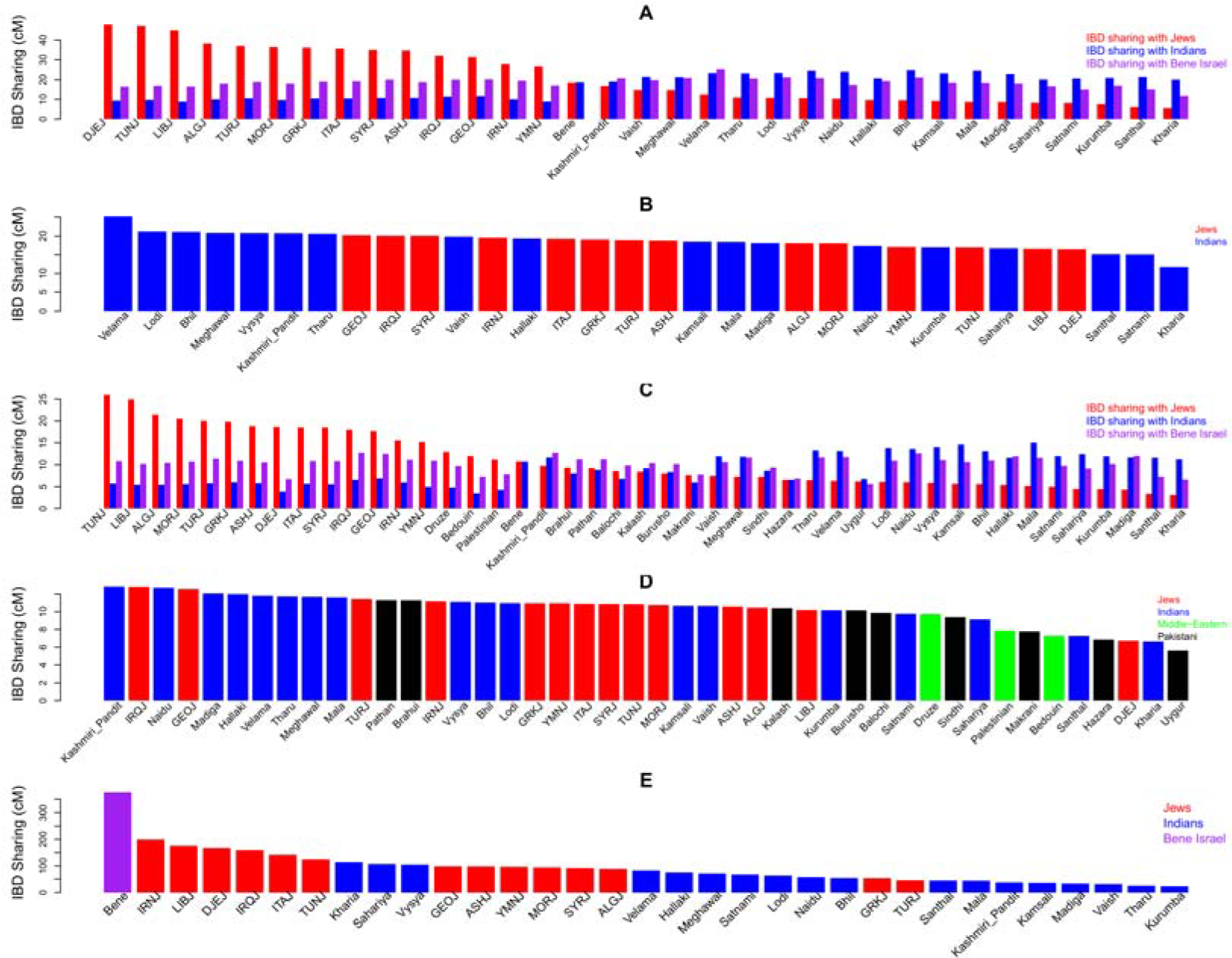
IBD sharing between and within Jewish, Indian, Pakistani and Middle-Eastern populations. (A) Average IBD sharing between different populations. For each population we measured its average IBD with Bene Israel (purple) and all other Jewish (red) and Indian (blue) populations. (B) IBD sharing of Bene Israel with other Jewish and Indian populations (C) Average IBD sharingbetween different populations. For each population from panel A, with the addition of Pakistani and Middle-Eastern populations, we measured its average IBD with Bene Israel (purple) and all other Jewish (red) and Indian (blue) populations. (D) IBD sharing of Bene Israel with other Jewish, Indian, Pakistani and non-Jewish Middle-Eastern populations. Analyses in panels C-D were performed on the dataset merged with HGDP dataset that contained smaller number of SNPs, and therefore the differences in IBD sharing values. (E) IBD sharing *within* populations.

Next, we used the merged dataset with Middle-Eastern and Pakistani populations for a lower-resolution IBD analysis. Still, Bene Israel showed higher IBD sharing with Middle-Eastern Jewish populations in this dataset as well (Figure 3C). Importantly, while the PCA, and ADMIXTURE showed that some Pakistani populations were more similar to Middle-Eastern and Jewish populations than Bene Israel, no Pakistani population showed as high IBD sharing with Jewish populations as compared to Bene Israel (Figure 3C). In addition, we used this merged dataset to examine whether the relatively high IBD sharing of Bene Israel with Jewish populations and mainly Middle-Eastern Jews is Jewish specific or Middle-Eastern in general. The IBD sharing of Bene Israel and non-Jewish Middle-Eastern populations was lower than their sharing with all other Jewish populations (Figure 3C; Figure S4B), implying that the link between Bene Israel and Jewish populations is at least in part Jewish-specific and not only Middle-Eastern. Although there were differences between the IBD sharing in the two datasets due to the different set of SNPs, there was overall significant correlation between the ranking of IBD sharing of Jewish (R=0.85, P-value=1.36e-4; Spearman correlation) and Indian (R=0.52, P-value=0.03, Spearman correlation) populations in the two datasets. Higher sharing with Middle-Eastern Jewish populations was also observed when we restricted the analysis to longer segments of IBD that reflect a more recent ancestor (Figure S5). While we did not see a clear trend for Indian populations showing higher IBD with Bene Israel, the higher-resolution analysis showed Velama to have the highest IBD sharing also in respect to longer IBD segments (Figure S5).

### Bene Israel as an admixed population of Jewish and Indian ancestral populations

Motivated by the above results, we next examined whether the Bene Israel community was an admixture of Indian and Jewish ancestral populations, using two different approaches as implemented in the ALDER [25] and GLOBETROTTER [26] tools. Given a putative admixed population and two populations that are taken as surrogates for the true ancestral populations, ALDER computes an admixture linkage disequilibrium (LD) statistic in the admixed population and uses it to examine whether the population is indeed an admixture of the ancestral populations [25]. In most cases of one Jewish and one Indian population taken as surrogate ancestral populations, there was a consistent and significant evidence for Bene Israel being admixture between these two populations (147 out of 252 possible pairs. Table S3 and Figure S6; the only population that did not show a significant evidence for being an ancestral population for Bene Israel was the Indian Kashmiri Pandit). Repeating the same analysis but with pairs of Indian populations or pairs of Jewish populations, as well as replacing Bene Israel with any other Indian or Jewish population did not find any pair with significant and consistent evidence for admixture, suggesting that the observed admixture for Bene Israel was not reflecting ANI-ASI admixture but a unique admixture between Jewish and Indian populations. ALDER admixture estimated time varied across the 147 significant pairs of populations, from ~19 (Iraqi Jews and Satnami) to ~33 (Georgian Jews and Mala) generations ago (650-1050 years ago, assuming 29 years per generation [18, 27]) with an average of ~25 generations (~820 years) ago (Table S2). These estimations place the admixture between a Jewish and Indian population well after the estimated time for the ANI-ASI admixtures of Indian populations (64-144 generations ago [18]) and after the establishment of many Jewish Diasporas [15] (Figure 4A). Turning to admixture proportions estimations based on ALDER, those estimated for Indian populations, varying between 44% (Vaish) and 20.2% (Kharia) were generally higher than that of Jewish populations, varying between 23% (Georgian Jews) and 15.5% (Libyan Jews; Figure 4B). When repeating ALDER analysis using the merged dataset with non-Jewish Middle-Eastern populations, the results were less significant, as expected by the smaller number of markers, but still many pairs of one Jewish/Middle-Eastern population and one Indian population were significant. Importantly, the results were more significant for Jewish populations as compared to non-Jewish Middle-Eastern populations: While 3 of the 17 (17.6%) Jewish/Middle-Eastern populations examined were non-Jewish, only 8 of the 113 (7.1%) significant pairs contained non-Jewish population, and all other pairs contained a Jewish population (Table S4 and Figure S7). Furthermore, when we replaced Bene Israel with Pakistani populations, there was no evidence for any Pakistani population being an admixed population with both Jewish and Indian ancestry. This result further emphasizes that the admixture detected by ALDER is not the ANI-ASI admixture.

**Figure 4.**
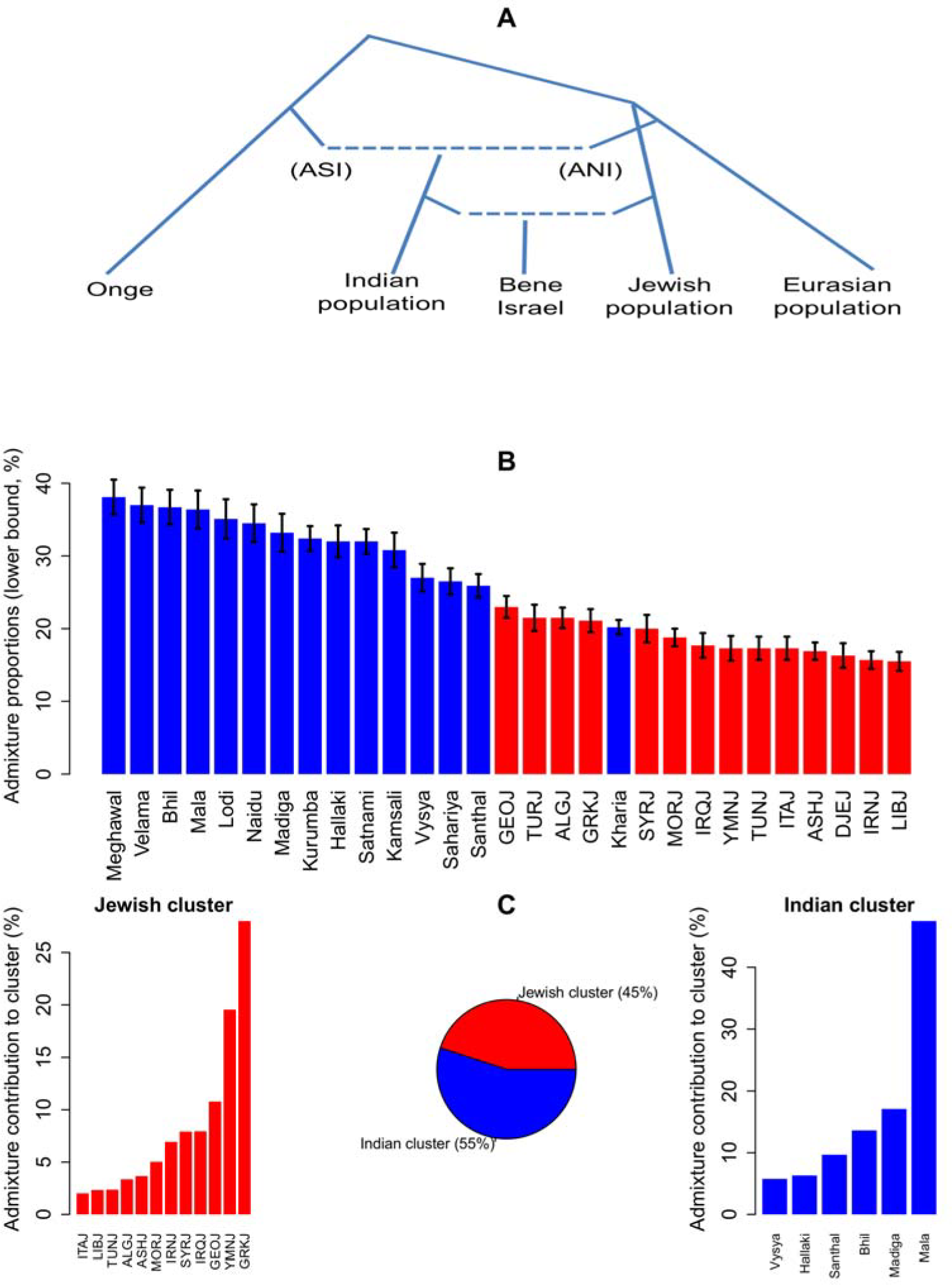
Bene Israel as an admixed population with Indian and Jewish ancestry. (A) Bene Israel as an admixed population of Indian and Jewish origin. Admixture time (~19-33 generations ago) is after the ANI-ASI admixture of Indian populations (64-144 generations ago). The ANI side is associated with Eurasian populations while ASI is associated with indigenous Andaman Island people (e.g., Onge). Dotted lines correspond to admixture between populations. This is a schematic overview of Bene Israel genetic history and branch lengths are not proportional to actual time. (B) ALDER admixture proportions estimations for Indian and Jewish populations being ancestral populations of the Bene Israel community. Values (with standard errors) are based on ALDER analysis with one-reference population. See also Tables S2 and S3. (C) GLOBETROTTER estimations for Indian (55%) and Jewish (45%) clusters proportions in Bene Israel admixture. The contribution of each population in each cluster is also presented.

In addition, we performed *f_4_*-based analysis [17] to test whether Bene Israel are closer to Jews than to non-Jewish Middle-Eastern populations (Materials and Methods). We found that Middle-Eastern Jewish populations were closer to Bene Israel as compared to other Middle-Eastern populations examined (Druze, Bedouin and Palestinians). Non-Middle-Eastern Jewish populations were still closer to Bene Israel as compared to Bedouin and Palestinians, but not as compared to Druze (Figure S8). These results further support the hypothesis that the non-Indian ancestry of Bene Israel is Jewish specific, likely from a Middle-Eastern Jewish population.

In addition, we also applied GLOBETROTTER on our dataset. GLOBETROTTER assigns haplotype segments of the admixed populations to different populations and uses the co-distribution of such segments from different populations to detect and infer admixture. In comparison to ALDER, which focuses on a pair of putative ancestral populations, it considers all populations and assigns ancestry component to all of them simultaneously. Importantly, GLOBETROTTER found evidence for admixture in the similar time range suggested by ALDER -27.7±0.43 generations ago. Admixture proportion estimation was 55% Indian and 45% Jewish. The main contribution from the Jewish cluster was from Greek, Yemenite and Middle-Eastern Jewish populations while the Indian cluster was mainly composed of populations with a high ASI component (Figure 4C). This may suggest that the ancestral Indian population had high ASI component. However, some of the ANI component of the ancestral Indian population may have been captured by the Jewish cluster, and resulting in the Indian side containing a higher ASI component. If the latter is true, GLOBETROTTER proportions estimation for the Jewish side (45%) is an overestimation of the true proportion as it also contains some of the ANI component from the Indian side.

We note here that GLOBETROTTER’s original study analyzed, among 95 worldwide populations, "Indian Jews" [26]. However, this group included members from both the Bene Israel and Cochin Jewish communities (four samples from each population [13]), and none of the Jewish populations examined here was used for the analysis. Nevertheless and reassuringly, they reported an admixture event occurring approximately 20 generations ago, with one side being Indian while the other side related to the Middle-East (e.g., South-Italians and Jordanians), which may partially reflect the admixture we report here for Bene Israel.

### High endogamy and founder event in Bene Israel

We now turn to examine the post-admixture population structure of Bene Israel. Both Jewish populations [14, 23], as well as Indian populations after the ANI-ASI admixture [17, 18], show high endogamy. We found that while Jewish populations showed higher IBD sharing as compared to Indian populations (P-value=4.84e-4, Wilcoxon rank sum test) the Bene Israel population exhibited a level that was almost as twice as high as any other of these populations (Figure 3D). Similarly, Bene Israel exhibits higher total length of homozygous segments (Figure 5A) and lower heterozygosity (Figure S9).

**Figure 5.**
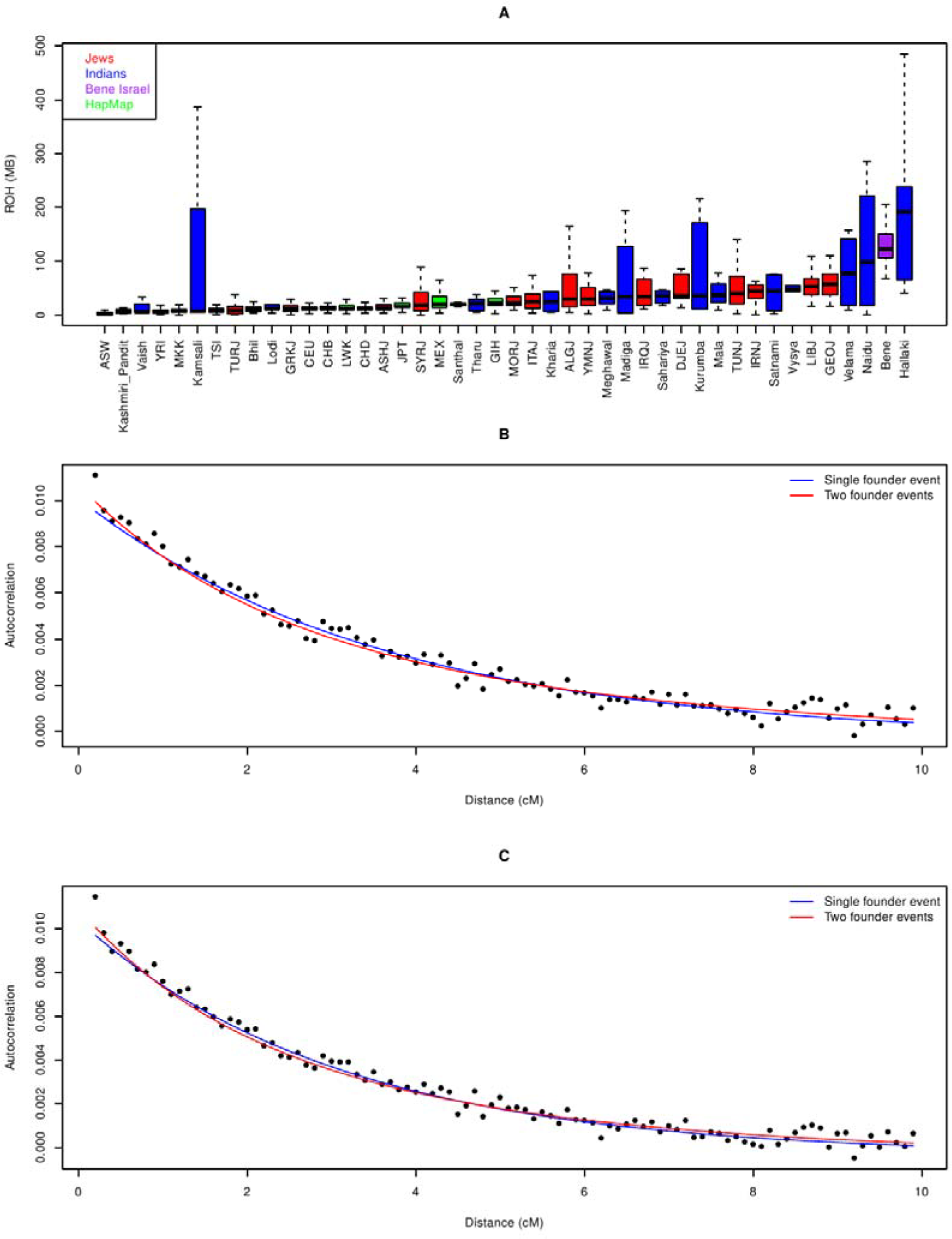
Founder events in Bene Israel population. (A) Total lengths of runs of homozygosity (ROH) in Jewish, Indian and HapMap populations. The larger variance in ROH values in some Indian populations is due to smaller sample size. (B-C) Autocorrelation in Bene Israel pairs, as a function of the genetic distance, after subtracting the autocorrelation between Bene Israel and other (B) Jewish and (C) Indian populations. Blue and red lines correspond to the fitted curve based on a single and two founder events, respectively.

These results can suggest not only endogamy but also a genetic bottleneck or a founder event where the contemporary Bene Israel population descended from a small number of ancestors. To directly examine this hypothesis, we used an allele-sharing statistic that measures the autocorrelation of allele sharing between individuals within a population and subtracts the cross-population autocorrelation to remove the ancestral autocorrelation effect. The decay of this statistic with genetic distance can verify if and when a founder event has happened [17, 28]. We applied this method to our dataset, using either all Jewish or all Indian populations for cross-population autocorrelation calculation and fitted it to one or two founder events (Figure 5B and Figure 5C). When fitted to a single founder event, analysis suggested it occurred 16 (using Jewish populations) and 14 generations ago (using Indian populations). Fitting to two founder events, which was slightly better, suggested a first founder event 30 generations ago followed by a second event 12 generations ago (using Jewish populations), and a founder event 26 generations ago followed by a second event 9 generations ago (using Indian populations). The first of these two events fits within the timescale of the admixture estimated above and may reflect the founding of this population in the admixing of Jews and Indians. The estimated time of 14-16 generations ago of a single founder event may be the average of these two founder events. The founder event and the genetic drift associated with it are reflected in several other results: by the relatively high F_ST_ values between Bene Israel and other Indian and Jewish populations (Figure S2) and by ADMIXTURE analysis which revealed that at K=8 Bene Israel form their own distinct cluster, though this population sample encompasses only 18 individuals out of 873 (Figure 2).

### Bene Israel admixture has been sex-biased

Lastly, we examined whether the ancestry of Bene Israel has been sex-biased using the Q ratio [29]. In a population with equal size of males and females, there are three copies of the X chromosome for every four copies of each autosome and therefore the expected genetic drift on the autosomes is 3/4 of the genetic drift on chromosome X, though this ratio is affected by many additional factors [29-31]. We found a significantly (P-value=5.11e-6, Wilcoxon rank sum test) lower ratio between Bene Israel and Jewish populations (Figure 6 and Table S5; mean=0.58) than between Bene Israel and Indian populations (Figure 6 and Table S5; mean=0.73; See also Figure S4). This entails that the Jewish contribution to Bene Israel has been smaller than otherwise expected for the X chromosome, which points to more male than female Jewish ancestors contributing to the formation of Bene Israel, consistent with findings from previous studies based on Y chromosomal analysis suggesting a paternal link between Bene Israel and Middle-Eastern populations [13]. As most of the Bene Israel samples in our dataset were women, we did not have enough power to analyze the Y chromosome, but mtDNA analysis revealed common Indian haplogroups (M, [32]), consistent with previous studies [8, 11, 13, 15] and the sex-bias we discovered above, while only a few samples had the H haplogroup which is common in Europe and in the Middle-East [33] but also present in lower frequencies in some Indian populations [34] (Table S6).

**Figure 6.**
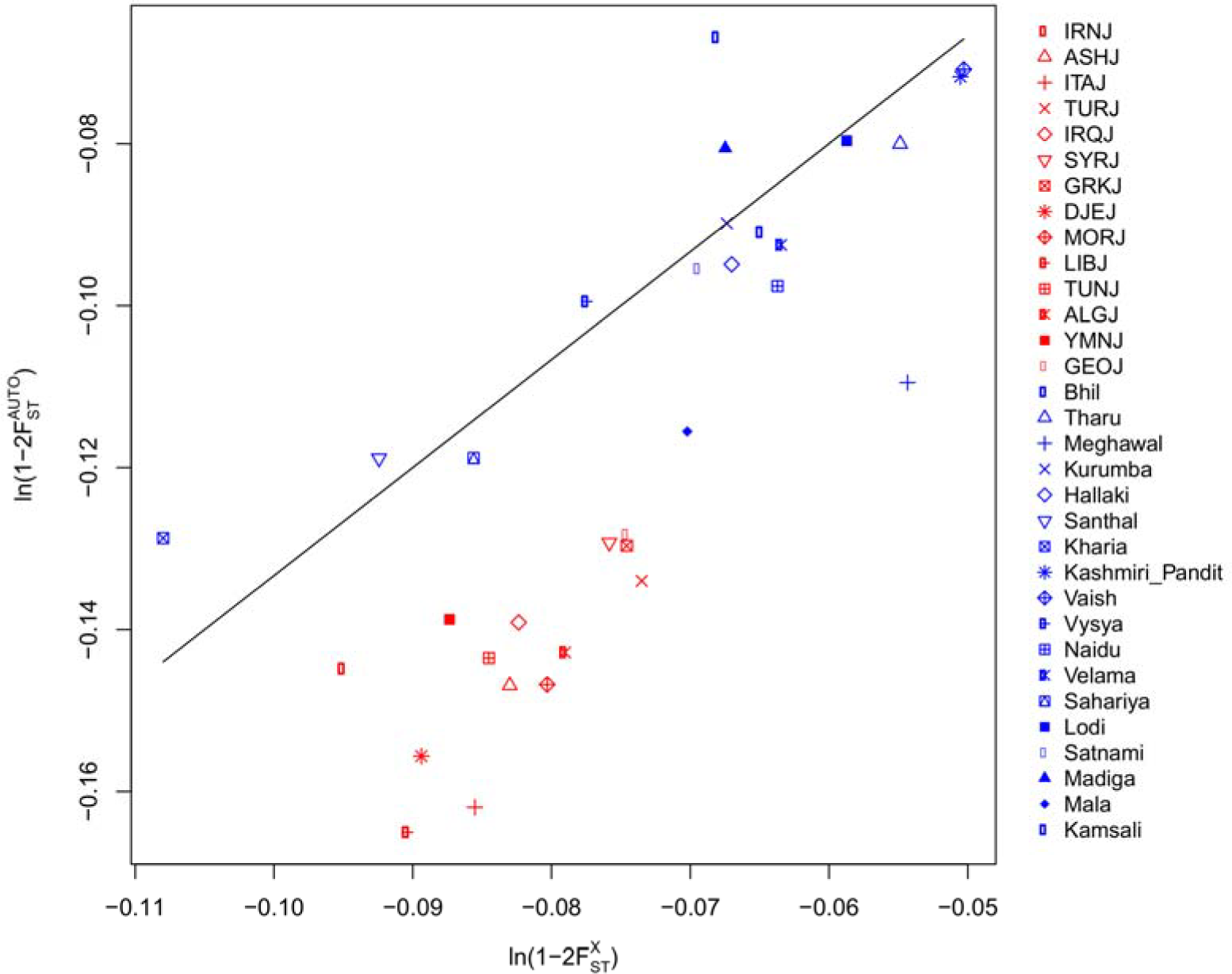
Autosome to chromosome X genetic drift ratio between Bene Israel and other populations. The line represents the ¾ ratio expected under the null hypothesis of similar demography of males and females in respect to the other populations examined (Jews and Indians, in our case). See also Table S5.

## Discussion

Previous studies used genetic markers to investigate the history of worldwide Jewish populations. While most Jewish Diaspora groups have been linked together and traced back to a Middle Eastern origin, in accordance with historical records [12-15], there were a few exceptions. Among these exceptions stood the Bene Israel community in India [15]. Autosomal markers failed to distinguish between Bene Israel and other Indian populations, with some suggestive evidence, based on uniparental markers, for a non-Indian component and a possible paternal link to the Middle East [3, 8, 11, 13, 15]. Furthermore, their history, beyond vague oral histories, remains largely unknown also in the presence of other, non-genetic, studies, highlighting the importance of a comprehensive genetic study of this population if we are to reveal their history. Indeed, a major advantage of this study over previous studies is the richness of the data. First, we obtained genome-wide genetic information for 18 Bene Israel samples as compared to only four samples in a previous genome-wide study [13]. In addition, the Indian dataset [17] was much more comprehensive as compared to previous studies on Bene Israel, containing genome-wide genotyping of 96 samples from 18 different Indian populations, in addition to 43 samples from 9 different Pakistani populations, some of the latter being also a result of an ANI-ASI admixture that gave rise to Indian populations [17, 18]. This detailed representation of the complex genetic history of India has been crucial in us addressing the challenge of inferring a unique Jewish contribution that is distinguishable from that of the ANI component in the ANI-ASI admixture [17, 18]. Furthermore, we were able to apply several recent tools that directly examine the hypothesis of admixture, and were not available when previous studies on Bene Israel were conducted. Our results partly support the oral history of Bene Israel by showing that the population is an admixed population, with the ancestral populations that contribute to the admixture being both related to Indian and to Jewish populations.

An important observation that sheds light on our results and their interpretation is the comparison between Bene Israel and Pakistani populations. PCA and ADMIXTURE results showed that compared to other Indian populations, Bene Israel were more similar to other Jewish and Middle-Eastern populations and showed much larger Middle-Eastern component than Indian populations. However, several Pakistani populations also showed similar trends and in some cases were even more similar to Jewish and Middle-Eastern populations as compared to Bene Israel. Therefore, although these results can be suggestive of a non-Indian genetic component in Bene Israel, they were not necessarily different from that of other Asian populations. However, when we applied several methods that are not based merely on allele frequencies, but rather consider the genetic distance between markers (e.g., ALDER) or long stretches of markers IBD that are shaped by more recent history, we observed clear evidence for a uniquely Jewish component in Bene Israel that is not shared by any other Indian or Pakistani population.

The unique Jewish component was observed by Bene Israel showing higher IBD sharing with Jewish populations as compared to all other Indian or Pakistani populations. In addition, while ALDER reported significant evidence for Bene Israel having Jewish and Indian ancestry, no other Indian or Pakistani population produced similar evidence for Jewish ancestry. Therefore, while the similarity between Pakistani and Jewish populations is likely a result of being both descendants of an ancestral Middle-Eastern population, we showed that the similarity between Jewish populations and Bene Israel is due to Bene Israel being direct descendants of Jews. If the link between Bene Israel and Jewish populations is more direct as compared to Pakistani population, why did some Pakistani populations seem to have more Jewish-related ancestry in PCA and ADMIXTURE? This is likely because Bene Israel is an admixed population, with only part of their ancestry being Jewish, while the other part is related to Indian ancestry. Indian populations, in turn, are more genetically distant from Jewish and Middle-Eastern populations as compared to the Pakistani populations under question.

By contrasting patterns on the X chromosome with the autosomes, we find that the admixture we detected was sex-biased, with relatively more Jewish males contributing to Bene Israel’s gene pool. This is in line with suggestive evidence based on mtDNA and Y chromosome [13]. Similarly, mtDNA analyses in this and previous studies [8, 11, 13, 15] show it is mainly of Indian origin pointed to a suggestive paternal link to Middle-Eastern [13]. It is the consideration of genome-wide data, though, that enabled us to examine and answer the question with accuracy.

The admixture we detected point to Bene Israel being admixed of both Indian and Middle-Eastern, likely Jewish, ancestry, with each of the two ancestries making a substantial contribution to its gene pool. The fact that the same analyses did not detect a similar trend to any of the other Indian or Pakistani population examined here, even those with relatively large ANI component, suggests that the admixture is not related to the ANI-ASI admixture. The uniqueness of the admixture event is further supported by its timing, which two distinct methods estimated consistently to be between 19 to 33 generations ago (~650-1050 years) ago (based on ALDER, while the point estimate based on GLOBETROTTER is 26 generations ago). Hence, this admixture event is much more recent than the ANI-ASI estimated admixture time (64-144 generations ago [18]).

Maimonides’ letter describing a Jewish population in India, which may be Bene Israel [4], was written ~800 years ago and is well within our estimated admixture time. This time is relatively recent as compared to Bene Israel oral history for the arrival of their Jewish ancestors to India (ranging from 8^th^ century BCE to 6^th^ century CE), yet none of these dates has any independent support [4]. Importantly, the admixture timing captures the timing of the actual interbreeding between Jewish and Indian populations, but it is plausible that Jews arrival to India predates the admixture. Similarly, our analysis assumes a single admixture event, but if several admixture events occurred, or if admixture has been more continuous, the estimated time of admixture may be intermediate between the different events, and biased towards the more recent admixture time [27].

While the admixture time is well after the ANI-ASI admixture and the forming of Jewish Diasporas, our analyses cannot suggest a unique pair of Indian and Jewish populations that are most likely to be the ancestral populations of Bene Israel, likely because of the large similarity between populations in each of these groups and the robustness of some of the methods (e.g., ALDER) when using proxy populations that did not directly descend from the true ancestral populations. Similarly, it is difficult to distinguish between Middle-Eastern and Jewish specific origin because of the Middle-Eastern origin of Jewish populations. However, our results suggest that the non-Indian component of Bene Israel ancestry is more likely to be Jewish-specific rather than Middle-Eastern more broadly, and that the Jewish forefathers of Bene Israel came to India from geographically close Middle-Eastern communities, perhaps through the Silk Road, and not from farther communities. Thus, Middle-Eastern Jewish populations showed larger IBD sharing with Bene Israel as compared to both other Jewish populations as well as other Middle-Eastern populations. Similarly, *f_4_*-based analysis suggested that Middle-Eastern Jewish populations are closer to Bene Israel as compared to non-Jewish Middle-Eastern populations. Finally, ALDER exhibited more significant results with Jewish populations as compared to non-Jewish Middle-Eastern populations. Regarding the ancestors from the Indian side, initial results based on PCA with both Indian and Jewish populations (Figure 1B) position Bene Israel close to Indian population with highest ANI ancestry. However, several analyses then suggest that this is likely due to the Jewish contribution to the population, and that the closest ancestral Indian populations are not necessarily those with the largest ANI component.

Further analysis of the genetic history of Bene Israel post-admixture reveals isolation and high endogamy in this population. While both Jewish [14, 15] and Indian populations [17] show high levels of endogamy, endogamy in Bene Israel is much higher as compared to these populations. Similar to many other Jewish Diasporas [15, 35], we find evidence for a founder event in Bene Israel. The estimated time of this event is ~14-16 generations ago, but we found evidence for two distinct founder events, the more recent one occurring more recently than these estimate and the more ancient one occurring at time of admixture. If indeed two founder events occurred, the estimated time of the first event provides further support to the admixture event and its timing, as admixture between a small group of Jews that arrived to India and local Indians is likely to be accompanied with a founder event, and indeed it is estimated to have occurred 26-30 generations (~850-970 years) ago. The isolation and the genetic drift experienced by Bene Israel have an effect on other types of analysis. For example, the 3-population test fails to find evidence for admixture by exhibiting positive *f_3_* values, likely due to the post-admixture genetic drift [17, 36], while the above methods that are less sensitive to such drift, e.g., ALDER [25], are able to detect it. The Bene Israel community was traditionally divided into two groups that in previous generations did not marry each other: Gora (or the “White” Bene Israel), presumed to be descendants of the seven couples who landed in the Konkan shore, and Kala (or the “Black” Bene Israel), presumed to be descendants of admixture between Bene Israel men and non-Bene Israel women [1, 2]. Our analysis did not provide evidence for the existence of two clear subgroups within Bene Israel samples. However, while the sample we analyzed consists of 18 Bene Israel individuals, it might still be too small or biased in its collection to detect both these subgroups.

Revealing the high endogamy and founder event(s) in Bene Israel is important not only from historical but also from medical perspective, as it predicts higher rates of recessive diseases within this population [37]. Indeed, a recent study on isolated foveal hypoplasia, a rare eye disease leading to poor vision, found that unrelated Bene Israel patients shared a homozygous mutation (c.95T<G, p.Ile32Ser) in the SLC38A8 gene [38]. Other recessive mutations in SLC38A8, a putative glutamine transporter, result in a similar medical condition [39]. The high prevalence of this mutation in Bene Israel (10% of Bene Israel individuals screened in the study [38]) which was completely absent from the entire set of individuals, including European and Indian, in the 1000 Genomes Project [40], reflects on the founder event and high endogamy this community has experienced.

A more complete history of Jewish Indian populations requires characterizing the history of Cochin Jews as well, which we pursued over the last few years in a separate study (see also Materials and Methods). The results are beyond the scope of the current study and will be published separately (Waldman, Biddanda, Dubrovsky, Campbell, Oddoux, Friedman, Atzmon, Halperin, Ostrer, Keinan; manuscript in preparation). They detail a complex history of Cochin Jews, with genetic contribution from both Indian and Jewish ancestry.

In conclusion, our results, based on an ensemble of different approaches, combine to support some versions of the oral history of the Bene Israel community as having both Jewish (likely from the Middle-East) and Indian origin, though the timing of this admixture is more recent as compared to their oral history. This study serves as an example of a genetic history study that does not only confirm known historical events but also reveal novel historical insights.

## Materials and Methods

### Recruitment of Bene Israel individuals

Recruitment of Bene Israel members occurred at Sheba Medical Center in Tel Hashomer, Israel (20 subjects) and at the Indian Bene Israel Community in Ramla, Israel (8 subjects) following approval of the study protocol and consent form by the Sheba Medical Center Helsinki Ethics Committee and the Director General of the Israeli Ministry of Health. All subjects provided written informed consent. The 20 samples collected at Sheba Medical Center were taken from individuals (who came for prenatal or oncogenetic counseling) identifying themselves as Indian Jews, rather than necessarily Bene Israel and therefore were either Cochin Jews or Bene Israel, the two large Indian Jewish communities in Israel. Although all individuals were recruited in Israel, where the vast majority of Indian Jews now live, they did not mix with non-Indian Jewish populations: all individuals recruited reported that their four grandparents belonged to the same Jewish community, similar to other Jewish populations analyzed in the current and in previous works [12, 14].

In order to distinguish between Bene Israel and Cochin Jews in the samples collected in Sheba Medical Center, we used the SMARTPCA tool from the EIGENSOFT software [41] and ADMIXTURE [22]. Briefly, we added to these 20 samples additional samples with known population of origin: 8 Bene Israel samples (from Ramla, as described above), 20 Cochin Jews samples (taken from the National Laboratory for the Genetics of Israeli Populations) and 36 Ashkenazi Jews samples [12]. Projection of these samples on the first two principal components (PCs) clearly showed three main clusters: Ashkenazi Jews, Cochin Jews and Bene Israel (Figure S9). Similarly, ADMIXTURE analysis (with K=3) suggested three main corresponding clusters (Figure S11). We labeled an Indian Jewish sample as either Bene Israel or Cochin Jewish if the ADMIXTURE estimated the fraction of the corresponding inferred cluster in this sample was at least 95% and if this was also visibly reflected in the PC analysis.

Following this procedure, eleven Indian Jews from Sheba Medical Center were labeled as Bene Israel. One of the Bene Israel samples was later removed in the quality control (QC) steps performed on the dataset, as described below, resulting in a total 18 Bene Israel samples used in the current analysis. In addition to that, we repeated PCA (Figure S12) and ADMIXTURE (Figure S13) analyses presented in the main text while using only the samples collected in Ramla and not those collected in Sheba Medical Center, obtaining similar results as those described in the main text for all Bene Israel individuals. Further support to our Bene Israel and Cochin Jews labeling was obtained from another independent source. Previously, Behar et al. [13] genotyped various worldwide Jewish populations, including Cochin Jews and Bene Israel (four members from each of these two populations). We merged our dataset with that dataset which was genotyped on a different platform (Illumina Human610-Quad v1.0 BeadChip), resulting in 50,483 shared SNPs. Projection of the first two PCs showed that our labeling was in accordance with the labeling of their samples (Figure S14). Cochin Jews have a different history [2], as also exhibited here in the above analyses and in previous genetic studies as well [11], and therefore they were not included in the current study. However, in parallel to this study, we pursued similar analyses to those described here to Cochin Jews samples. The results of these analyses will be published separately (Waldman, Biddanda, Dubrovsky, Campbell, Oddoux, Friedman, Atzmon, Halperin, Ostrer, Keinan; manuscript in preparation).

### Jewish dataset and genotyping

We included in this study a Jewish dataset containing samples from additional 14 Jewish populations, which were collected as described previously [12, 14] (Table S1). All samples, including the Bene Israel samples in this study, were genotyped on the Affymetrix 6.0 array (Affy v6) at the genomic facility at Albert Einstein College of Medicine. Compared to previous studies [12, 14], we used an updated version (1.5.5) of the Birdseed tool in the Birdsuite software [42] to recall the genotypes again. Samples with ambiguous gender (based on genotyping) were removed from the analysis. For the current study, where we focused on Bene Israel, we did not include Ethiopian Jews as previous studies have shown that they do not group tightly with other Jewish populations [13, 14].

### Indian dataset

We also incorporated a dataset of Indian populations based on the study of Reich et al. [17] where the samples were genotyped in the same array (Affymetrix 6.0). We removed Indian populations living in islands and not the mainland (Great Andamanese and Onge), and those defined by Reich et al. [17] as genetic outliers (Aonaga, Nysha and Siddi, Chenchu). In addition, we removed Srivastava from our analysis, as after QC steps only one sample from this population was left, resulting in the inclusion of 18 Indian populations in our analysis (Table S1). The dataset generated by Reich et al. [17] contained, in addition to the Indian populations, samples from HapMap3 [43] populations and these samples were also used (after QC, described below) for phasing and for some of the analyses (Table S1).

### Human Genome Diversity Project dataset

For some of the analyses we also included data for non-Jewish Middle-Eastern populations (Bedouin, Druze and Palestinian), and nine different Pakistani populations (Kalash, Balochi, Brahui, Makrani, Sindhi, Pathan, Burusho, Hazara and Uygur). Middle-Eastern populations were used to distinguish between Middle-Eastern and Jewish specific genetic attributes while Pakistani populations were taken as populations with high ANI component that reside geographically between India and the Middle-East. We incorporated these samples by merging our dataset with data from the Human Genome Diversity Project (HGDP) [44] genotyped on the Affymetrix GeneChip Human Mapping 500K [19]. This dataset included five unrelated samples from each of these populations, except Makrani and Sindhi with four samples each that passed QC steps (Table S1).

### Dataset merging and Quality Control

After removing SNPs with low call rate (<95%) from the two datasets (Jewish and Indian), we merged them together (including the HapMap samples in the Indian dataset [17]) and removed individuals as follows:

1. Relatives. Following Campbell et al. [14], which analyzed most of the Jewish populations present in the current study, two individuals were considered related if their total autosomal identity by descent (IBD) sharing was larger than 800 cM and if they shared at least 10 segments with length of at least 10 cM (see below how IBD sharing was calculated). To remove as few related individuals as possible while maintaining only unrelated individuals in our dataset, we constructed a graph whose vertices were individuals, and two individuals were connected if they were defined (according to the above criteria) as related. We then tried to find a maximal independent set (i.e., a maximal set of unrelated individuals) in this graph using a greedy algorithm [45].
2. Genetic outliers. We used the SMARTPCA program [41] to detect genetic outliers, with default parameters for genetic outlier removal. We removed individuals further than six standard deviations from the mean in any of the top ten eigenvectors over five iterations. This analysis was done for each population alone, based on autosomal SNPs.

The merged dataset following these QC steps included 513,581 and 25,379 autosomal and X chromosome (in the non-pseudoautosomal regions) single nucleotide polymorphisms (SNPs), respectively, for 461 individuals from 33 Jewish and Indian groups. Additional 844 samples from 11 HapMap3 populations were also available in this dataset, resulting in total 1305 samples. Further merging with the HGDP dataset (for some analyses) consists of 1363 samples with 304,973 shared autosomal SNPs. The number of samples from each population is shown in Table S1.

A set of filtered SNPs based on linkage disequilibrium (LD) was used in the following analyses: PCA, F_ST_, ADMIXTURE, runs-of-homozygosity and heterozygosity. For each pair of SNPs showing LD of r^2^>0.5 we considered only one representative (using SMARTPCA’s [41] r2thresh and killr2 flags). This filtering was done for each analysis alone, depending on LD in the specific set of populations used in the analysis. Other analyses were performed on the full datasets described above.

### Identity-by-descent analysis

We phased the data with the BEAGLE software (version 3.3.2) [46] and extracted shared identity-by-descent (IBD) segments with GERMLINE (version 1.51) [24], using the same parameters described in Campbell et al. [14] which analyzed most of the Jewish populations examined here. To reduce the rate of false positive IBD segments, only segments with length of at least 3 cMs were considered for analysis. Similar to previous studies [12, 23], we ignored regions with low informative content. Specifically, using non-overlapping windows (of 1 MB or 1 cM) we ignored all regions with SNP density of less than 100 SNPs per cM or per MB. Genetic positions were obtained from the HapMap genetic map (downloaded from: ftp://ftp.ncbi.nlm.nih.gov/hapmap/recombination/2011-01_phaseII_B37/).

For each pair of unrelated individuals we calculated the total length of autosomal IBD sharing. Given two populations, the average IBD sharing of these two populations was defined as the average IBD sharing between all pairs of individuals from these populations. Similarly, the average IBD sharing within a population was defined as the average IBD sharing between all pairs from this population. In addition, we also calculated the average IBD sharing between a population and the group of all Jewish populations, by averaging the IBD sharing between this population and each of the other Jewish populations. We repeated a similar procedure to calculate average IBD sharing between a population and the group of all Indian populations. As this analysis was done to compare other populations to Bene Israel, the Bene Israel population was not considered to be either Jewish or Indian in this analysis.

We obtained empirical estimations for the distribution of average IBD sharing between populations by sampling 10,000 times ten Bene Israel individuals and measuring the average IBD between these individuals and members of other populations. These distributions were used to obtain standard error estimates for the average IBD sharing between Bene Israel and these populations. In addition, we used these distributions to examine whether Middle-Eastern Jewish populations show higher IBD sharing with Bene Israel as compared to other Jewish populations. This was done by comparing the average IBD sharing distributions of two Jewish populations (one of them being Georgian, Iraqi, Syrian or Iranian Jews) using Wilcoxon rank sum test.

### Principal Component Analysis

Principal component analysis (PCA) was performed using the SMARTPCA program [41] on the Jewish, Indian, and several additional worldwide populations. As the number of samples from each population can affect the results of PCA [20] and as there were more samples in each of the Jewish populations as compared to each of the Indian populations, we repeated PCA using not more than four samples from each population selected randomly (using the popsizelimit flag in SMARTPCA).

### F_ST_

We calculated population differentiation based on differences in allele frequencies between each pair of populations using the F_ST_ statistic, following the definitions described previously in Reich et al. [17]. Specifically, let *p_i_* be the frequency of a variant in a biallelic SNP in two populations (*i* = 1,2) and define *q_i_* = 1 – *p_i_*. We defined F_ST_ as

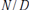

where

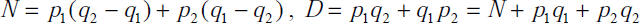

To generalize this measure to more than a single SNP, we followed the “ratio of averages” approach [47], where *N* and *D* were averaged separately and only then their ratio was taken. Thus, let *N_i_*, *D_i_* be the above definitions for SNP *i*, then for a set of SNPs *S*, F_ST_ was defined as:

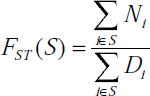

Similar to Reich et al. [17], the following estimators for *N* and *D* were used:

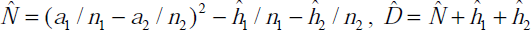

Where *a_i_* and *b_i_* are the allele counts of the two alleles, and *n_i_* = *a_i_* + *b_i_*. We calculated F_ST_ separately for the autosomes and for the X chromosome.

## ADMIXTURE

ADMIXTURE [22], a STRUCTURE [48] like algorithm, assigns for each individual its proportion in any of K hypothetical ancestral populations, and therefore can reveal relations between different populations. ADMIXTURE (version 1.2) analysis was performed with default parameters and varying values of K (from K=3 to K=10), with 200 bootstrap replicates. We ran ADMIXTURE with the dataset merged with the HGDP populations and included, in addition to the 33 Jewish and Indian populations, the following populations: CEU, YRI, CHB, JPT Druze, Bedouin and Palestinians, and nine Pakistani populations (total 873 unrelated samples). The dataset was filtered based on LD, resulting in 174,421 autosomal SNPs. ADMIXTURE’s cross validation procedure was used to determine the K that fits the data best.

### Inferring admixture proportions and time

We applied two tools to examine the hypothesis that Bene Israel population was an admixed population and to infer admixture proportions and time: ALDER (version 1.03) [25] (version 1.03) and GLOBETROTTER (downloaded in March 2015) [26]. A detailed description is found in the original publications of these tools, while we provide a brief description in the following.

(1) ALDER: Given a putative admixed population and two surrogate populations taken as a proxy for the presumed ancestral populations, ALDER uses admixture LD statistic to look for evidence for admixture (assuming a single admixture event). For each pair of SNPs ALDER calculates a statistic being the covariance of these two SNPs in the admixed population, weighted by the allele frequency differences between the two reference populations. Exploring the behavior of this admixture LD statistic as a function of the genetic distance between the two SNPs can imply whether the population is admixed or not.

ALDER fits the statistics curve to an exponential function

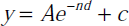

where *n* is the number of generations since admixture and *d* is the genetic distance (in Morgans). In addition to the test of admixture using two reference populations, ALDER examines evidence for admixture using only one surrogate population as a reference, with the admixed population serving as a proxy for the second population.

We considered a pair of populations as a candidate for being the ancestral populations for a certain population if all three ALDER results (two one-reference admixture LD and a two-reference admixture LD analyses) were significant and the estimated time of decay was consistent between the three.

In addition to time of admixture, ALDER also estimates admixture proportions from the amplitude of the exponential curve. This is done both in the one-reference version of ALDER (estimating the lower bound of admixture proportion of that population) as well as in the two-reference version. As the populations examined here are taken as a proxy for the true mixing populations, the admixture proportions suggested are lower bounds [25]. A caveat in the two-reference version admixture proportion estimation is that ALDER does not determine to which population to assign the admixture proportion estimation α (i.e., it does not distinguish between α and 1 – α). Therefore, we used min(α,1 – α) as a lower bound for the admixture proportion of the Jewish population in each significant pair. To determine α from the output of ALDER two-reference population test, *f*_2_ values (representing genetic drift between the two populations [17]) are needed [25] and these were calculated using MixMapper [49].

(2) GLOBETROTTER: Given a putative admixed population and a set of populations (some of them may be a proxy for the presumed ancestral populations), GLOBETROTTER examines whether the putative admixed population is an admixed population of some of the populations from that set [26]. As GLOBETROTTER is based on haplotypes, we phased the data using BEAGLE [46]. GLOBETROTTER algorithm requires several steps. First, the chromosomes of each individual in the admixed population are broken into ‘chunks’ where each chunk is assigned, based on similarity, to a single individual from one of the other populations. This step, implemented in the CHROMOPAINTER [50] tool results in ‘coloring’ of the chromosomes of admixed individuals with different populations. Second, for each pair of populations a curve, which quantifies each genetic distance how often a pair of haplotype chunks separated by this distance come from each pair of populations, is produced. Similar to ALDER, the decay rates of these curves are used to examine whether admixture event happened and to infer its time, while the amplitude of the curve is used to infer the contributing populations and their proportions. In case of evidence for admixture, GLOBETROTTER also examines whether the data fits better single exponential decay (i.e., single admixture event) or a mixture of exponential decays (i.e., several admixture events or continuous admixture over a longer period). In case of admixture, GLOBETROTTER suggests two main clusters of admixture, each may be composed of several populations, which together represent the genetic structure of the ancestral population.

We used CHROMOPAINTER (version 2) and ran GLOBETROTTER on Bene Israel and the Jewish and Indian populations, using 100 bootstrap replicates to obtain standard error estimates for the admixture time.

### Time estimates

We converted the number of generations into years by assuming 29 years per generation for such recent history [18, 27] and that individuals genotyped in the current study were born circa 1950 CE. Thus, if *n* is the number of generations since admixture, we convert it to the year 1950 – 29(*n* + 1) (CE). Changes in generation lengths estimations will scale the time estimations proportionally.

### Homozygosity and Heterozygosity estimations

We used PLINK (version 1.07) [51] to identify runs-of-homozygosity (ROH) –autozygous segments in the genome. We used the following flags in PLINK: ‘--homozyg --homozyg-window-kb 1000 --homozyg-window-snp 100 --homozyg-window-het 1 --homozyg-window-missing 5 --homozyg-snp 100 --homozyg-kb 1000’.

The heterozygosity score of an individual was defined as the fraction of the heterozygous SNPs among all autosomal SNPs (after LD-pruning).

### Estimating founder event time

We used allele sharing autocorrelation for estimating time of founder event, along the lines suggested by recent studies [17, 28]. Specifically, for each pair of individuals from the population, and for each autosomal SNP, we measure the number of alleles these individuals share: zero, one or two. When both of the individuals are heterozygous for the SNP, we consider them as sharing one allele (to account for haplotype phasing ambiguity). Thus, each SNP is represented by a vector where each entry in the vector corresponds to a pair of individuals and the value of that entry is the number of shared alleles between these two individuals. Next, a Pearson correlation coefficient is calculated between the vectors for each pairs of SNPs (referred as allele sharing autocorrelation). To remove the effect of ancestral allele sharing autocorrelation, we subtract the cross-population allele sharing using this population and a different population. To infer the founder event, we plot the autocorrelation vs. genetic distance and fit the curve to the exponential equation

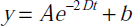

where *t* represents the number of generations since the founder event and *D* is the genetic distance (in Morgans) between the two SNPs [17, 28].

We applied this method for Bene Israel and calculated allele sharing autocorrelation between each pair of SNPs less than 30 cM apart. We partitioned the values into 0.1 cM bins and considered the mean of each bin. To consider two founder events, we fitted the decay to an equation of the form

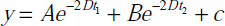

where *t_1_* and *t_2_* were the times (in generations) since the two founder events. Fitting was done by non-linear least squares, using the *nls* function in **R** [52]. Evaluation between the single and two founder events models was done by comparing the sum of residuals of each of the models.

### Sex-biased population differentiation

To examine sex-biased demography, we calculated a statistic presented by Keinan et al. [29]: It estimates differentiation in allele frequencies (measured by F_ST_) between two populations for the autosomes 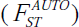 and for the X chromosome 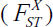 to estimate a ratio

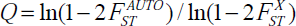

*Q* captures the relative genetic drift between the X chromosome and the autosomes. Under several assumptions [29], if the effective population size of males and females has been equal since the two populations split, *Q* is expected to be ¾, the ratio of effective population size of the X chromosome to the autosomes in this case. A significant deviation from 3/4 may suggest sex-biased demography since population split.

### mtDNA analysis

mtDNA genotypes were used to assign to each of the Bene Israel samples a mtDNA haplogroup basedon HaploGrep classification [53].

### *f_4_* analysis

We performed f4 test [17, 36] of the type *f_4_*(YRI, Bene Israel; J, ME) where J and ME are Jewish and Middle-Eastern populations, respectively. Assuming YRI is an outgroup for all other populations (Bene Israel, J and ME), If ME is an outgroup in respect to Bene Israel and J,*f_4_*(YRI, Bene Israel; J, ME) is expected to be negative, while if J is an outgroup in respect to Bene Israel and ME, it is expected to be positive [17, 36]. For the analysis we also assume that the Indian ancestry of Bene Israel does not affect the results (i.e., the split between the Indian ancestors of Bene Israel and Jewish populations occurred before the split between Jewish and Middle-Eastern populations).

## Acknowledgments

We thank all the individuals who contributed DNA for this study. We thank David Reich for providing us with the Indian dataset. We thank Po-Ru Loh and Mark Lipson for providing us with ALDER, advice on how to use it and interpret its results, and for comments on our results. We thank Garrett Hellenthal for providing us with GLOBETROTTER and advice on how to use it. We also thank Priyanka Vijay and Eyal Nitzany for preliminary analysis of related data, and Amy Williams and members of the Keinan and Halperin labs for helpful discussions. We also thank two anonymous reviewers for their comments.

AK and YYW were supported in part by National Institutes of Health Grant R01HG006849. AK was also supported in part by National Institutes of Health Grant R01GM108805 and by The Ellison Medical Foundation and the Edward Masllinckrodt, Jr. Foundation. EH is a faculty fellow of the Edmond J. Safra Center for Bioinformatics at Tel Aviv University. EH and YYW were partially supported by the Israeli Science Foundation (grant 1425/13), by the United States-Israel Binational Science Foundation (Grant 2012304) and by the German-Israeli Foundation (Grant 1094-33.2/2010). YYW was also partially supported by the Len Blavatnik and the Blavatnik Family Foundation. NRD was supported by the Tri-Institutional Training Program in Computational Biology and Medicine (via National Institutes of Health training grant 1T32GM083937). PBR was supported by the National Science Foundation Graduate Research Fellowship (grant No. 2013172342).

